# A neural network-based method for exhaustive cell label assignment using single cell RNA-seq data

**DOI:** 10.1101/2021.09.25.461825

**Authors:** Ziyi Li, Hao Feng

## Abstract

The fast-advancing single cell RNA sequencing (scRNA-seq) technology enables researchers to study the transcriptome of heterogeneous tissues at a single cell level. The initial important step of analyzing scRNA-seq data is usually to accurately annotate cells. The traditional approach of annotating cell types based on unsupervised clustering and marker genes is time-consuming and laborious. Taking advantage of the numerous existing scRNA-seq databases, many supervised label assignment methods have been developed. One feature that many label assignment methods shares is to label cells with low confidence as “unassigned.” These unassigned cells can be the result of assignment difficulties due to highly similar cell types or caused by the presence of unknown cell types. However, when unknown cell types are not expected, existing methods still label a considerable number of cells as unassigned, which is not desirable. In this work, we develop a neural network-based cell annotation method called NeuCA (Neural network-based Cell Annotation) for scRNA-seq data obtained from well-studied tissues. NeuCA can utilize the hierarchical structure information of the cell types to improve the annotation accuracy, which is especially helpful when data contain closely correlated cell types. We show that NeuCA can achieve more accurate cell annotation results compared with existing methods. Additionally, the applications on eight real datasets show that NeuCA has stable performance for intra- and inter-study annotation, as well as cross-condition annotation. NeuCA is freely available as an R/Bioconductor package at https://bioconductor.org/packages/NeuCA.

## Introduction

Single-cell RNA sequencing (scRNA-seq) provides an unprecedented ability to characterize the transcriptome heterogeneity at the single cell level. This technology has been applied in many research areas including evolutionary^1^, immunology^2^, neurodegenerative diseases^3^, and cancer^4,5^. With the fast-evolving technology and the growing interests of scRNA-seq applications, a vast number of datasets have been collected and annotated by domain experts in the past decade. These data deepened our understanding of the intrinsic characteristics of transcriptomics, as well as the phenotypes associated. Simultaneously, there also are community efforts that work towards a comprehensive collection of single cell profiling, such as the Human Cell Atlas^6–8^ and Mouse Cell Atlas^9^. From these joint efforts, massive amounts of scRNA-seq data were generated to provide excellent cell type references for understanding and annotating new data.

Because the sequencing output of scRNA-seq is anonymous in terms of cell identities, annotating the sequenced cells is usually the initial step in single cell data analysis. Traditionally, researchers adopted a two-step approach: first, applying unsupervised methods to cluster the cells; second, identifying the cell labels using known cell type markers. Many single cell clustering methods were proposed, including Seurat^10^, SC3^11^, TSCAN^12^. However, this two-step approach is a time-consuming and laborious procedure, which becomes even more challenging if good marker genes are unavailable to researchers. Recognizing the limitations, a number of supervised approaches were proposed to utilize existing single cell datasets and directly annotate cells without the clustering step. Popular methods in this category include scmap^13^, CHETAH^14^, CellAssign^15^, and others. In general, if the tissue type of interest has never been studied before, there will be no existing dataset available for training supervised methods. An unsupervised method becomes the only viable solution in such circumstance. However, when training data are available, the improved accuracy, speed, and reproducibility of supervised methods easily outperform unsupervised ones.

Supervised methods can be categorized into three different groups: marker based, correlation based, and tree structure based. Marker-based methods, as the name indicates, assign cell labels based on the provided marker genes^15,16^. Methods in this category require known knowledge of cell type-specific marker genes, and they thereby could be prone to marker misspecification. Correlation-based methods assign cell labels by cell-to-cell correlations or other similarity metrics^13,17^. For cell types that are closely correlated, the pairwise correlations of several cell types could be high simultaneously. The methods in this category may have difficulty accurately assigning labels under these high correlation settings. Tree structure based methods take the advantage of tree structures of cell types. This method type selects different sets of features at each classification bifurcation, which coincides with the idea of several recent publications that the cell types in a tissue form a hierarchical structure^18–20^. Such information could be used to improve classification accuracy of closely correlated cell types. For example, A popular use of this approach is the CHETAH method^14^. It is specifically designed for the situation when unknown cell types (e.g., tumor cells) present in the data. However, when researchers expect a tiny portion or no unknown cell types from a dataset of interest, CHETAH still produces an unignorable amount of unassigned/intermediate cell labels, as shown in our simulation study and real data analysis later.

Recognizing the limitations of existing methods, we developed a new method for exhaustively annotating cells in scRNA-seq data. We utilized well-studied tissues from existing scRNA-seq datasets with known labels as the training data. Of note, the training data have covered most cell types, thus few if any unknown cell types are expected. This assumption generally holds for many tissue types, including human brain^3^, human pancreas^21^, mouse brain^22^, mouse retina^23^, and many other tissue types under normal conditions^24^. When closely correlated cell types exist, our approach uses the tree structure of the cell types through a hierarchy of neural networks to improve annotation accuracy. At the same time, feature selection is performed in the hierarchical structure to further improve classification accuracy. In situations where cell type correlations are not high, we adopt a feed-forward neural network. We call our whole pipeline the <u>Neu</u>ral network-based <u>C</u>ell <u>A</u>nnotation (NeuCA). Leveraging well-understood tissues, NeuCA annotates cells in an exhaustive way to avoid loss of cells for downstream analysis.

Through a series of simulations based on real data, we comprehensively evaluated NeuCA in different settings and compared its performance with five popular existing supervised/unsupervised methods. Our applications on eight real datasets showed that NeuCA maintained stable performance in inter- and intra-data annotation, as well as cross-condition annotation. Our proposed method NeuCA is available through a freely available R/Bioconductor package at https://github.com/haoharryfeng/NeuCA.

## Results

### Method overview

NeuCA is a supervised cell label assignment method. It uses existing scRNA-seq data with known labels to train a neural network-based classifier and then predict cell labels in new data of interest. Figure 1 provides a schematic overview of the proposed method. Based on the training data, NeuCA first obtains the mean gene expression profile for each cell type and calculates the correlation matrix across cell types. NeuCA then will choose one out of the two approaches as described. If the correlation matrix contains highly correlated cell types (defined as cell types with Pearson’s correlation coefficient ≥ *τ*), NeuCA constructs a tree structure by hierarchical clustering and trains a series of neural networks based on this tree structure. We used *τ* = 0.95 for the following experiments, but users can specify this value in the R package. Predicted labels are obtained by applying the trained hierarchical neural network model to the testing data. In constructing the hierarchical neural network, different sets of features(genes) are selected at each iteration to accommodate the differences between cell types. If the correlation matrix does not show the existence of highly correlated cell types, NeuCA will train a feed-forward neural network for cell type prediction. This simplified approach is motivated by the fact that a tree structure is unnecessarily expensive when cell types are not similar. In this case, a feed-forward neural network can achieve satisfactory prediction accuracy. Collectively, NeuCA calculates the correlation matrix first and determines its adopted approach from the two strategies described above.

**Figure 1.**
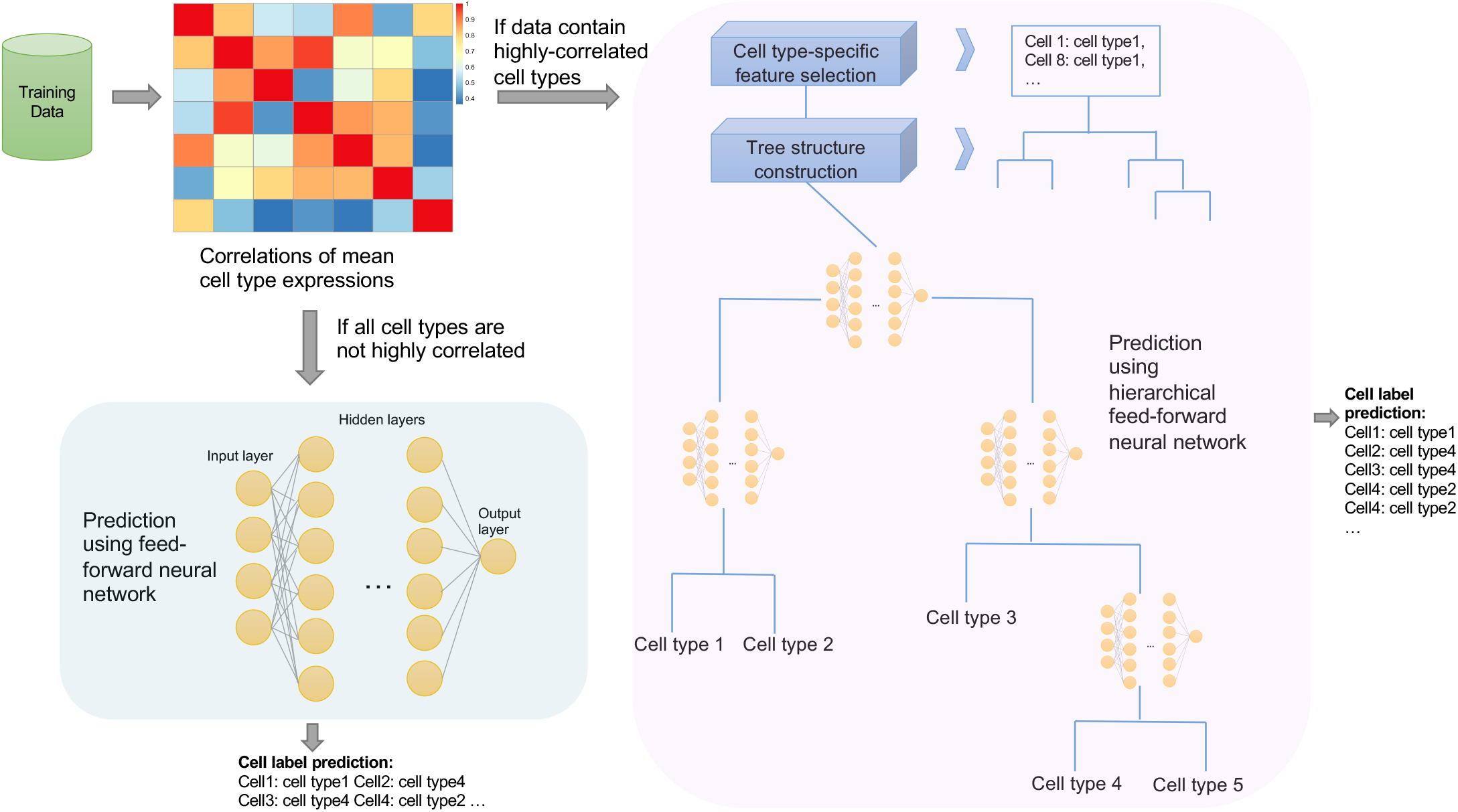
A schematic overview of the proposed method. Based on the correlation matrix of the training data, NeuCA will detect if highly correlated cell types exist, and decide between the following two routes: (1) with the presence of highly correlated cell types, NeuCA will adopt a hierarchical model with neural networks for cell label identification (right, pink panel); (2) in the absence of highly correlated cell types, a feed-forward neural network will be adopted for cell label identification (lower-left, cyan panel).

The advantages of the proposed method are threefold. First, NeuCA can recognize whether the cell types are highly correlated or not, and adjust its strategy. The presence of highly correlated cell types is usually an indicator for the increased complexity of the prediction task. Under an increased difficulty level, the combined approach of a tree structure and neural network model will show its merits. Under the scenario where high correlation is absent, the feed-forward neural network usually works well. Second, the adoption of a cell type hierarchical tree and stepwise feature selection in NeuCA help further improve accuracy. The hierarchical structures of cell types have been identified and used in several previous works^14,19,25^ for their advantage in analyzing cell types with high similarities. When numerous cell types are available, the hierarchical approach of cell annotation disassembles a complicated problem to a series of simple binary classification problems. For example, among peripheral blood mononuclear cells, it is easier to distinguish CD4T cells from CD8T cells, given only the mixture of these two cell types, than mixing them with all other peripheral blood cell types. In addition to hierarchical classification, we select the most distinguishable features at each step, which further improves similar cell type separation accuracy. Third, NeuCA assignments are exhaustive, in that they do not allow cells to stagnate at the intermediate node of the tree. This idea is supported by the assumption that most or all cell types in the new testing data have been included and correctly labeled in the training data. This eliminates the unnecessarily high unassignment rate in well-studied tissues. In the following numerical and real data experiments, we showed this would improve assignment accuracy.

### Numerical experiments with cell-sorted PBMC data

We first applied our proposed method, NeuCA, and other existing methods on a series of peripheral blood mononuclear cell (PBMC) datasets. All the experimental datasets generated in this section are based on the 10X PBMC scRNA-seq data^26^. This dataset contains single cell transcriptome data of more than 60,000 cells from fluorescence-activated cell sorting (FACS). The FACS experiment provided the gold standard of cell labels for all the sequenced cells, making it a reliable resource for benchmarking. We designed a series of experiments with randomly drawn cells from this dataset to serve as Monte Carlo simulation studies. This allowed us to fully evaluate the proposed and existing methods under various scenarios.

The methods benchmarked here included three supervised cell annotation methods, NeuCA, scmap, and CHETAH, and two unsupervised clustering methods, Seurat and SC3. We tested NeuCA with three different numbers of nodes: relatively large, medium, and small (see the Method section for details), denoted as NeuCA-big, NeuCA-med, and NeuCA-small, respectively. We also compared two versions of scmap, scmap-cluster, and scmap-cell, representing cluster-wise and cell-wise approaches for cell annotation. The two evaluation metrics were accurately assigned rate, calculated as the proportion of correctly classified cells over total cells, and adjusted Rand index (ARI), an indicator of the similarities between two clustering results. For unsupervised methods, we reported ARI only since matched cell labels are not directly available.

### Overall comparisons

We provided an overall comparison between NeuCA and existing methods across various scenarios. These scenarios included various training sets using 10%, 20%, 50%, or 80% of all PBMC data and the testing sets using 800, 1600, or 4000 randomly selected PBMC cells. Figure 2A and 2B present the overall accurately assigned rate and ARI of the evaluated methods. All three versions of NeuCA ranked at the top, indicating high overall accuracy and high consistency with the truth. With NeuCA-med, we achieved a 10% improvement in accuracy, compared with the best existing approach, scmap-cluster. We achieved at least 5% accuracy gain compared with scmap-cluster, even with NeuCA-small. Results of ARI indicate a highest concordance between our predicted cell clustering labels and the true labels. These results show that NeuCA outperforms both unsupervised and supervised methods that are currently available.

**Figure 2.**
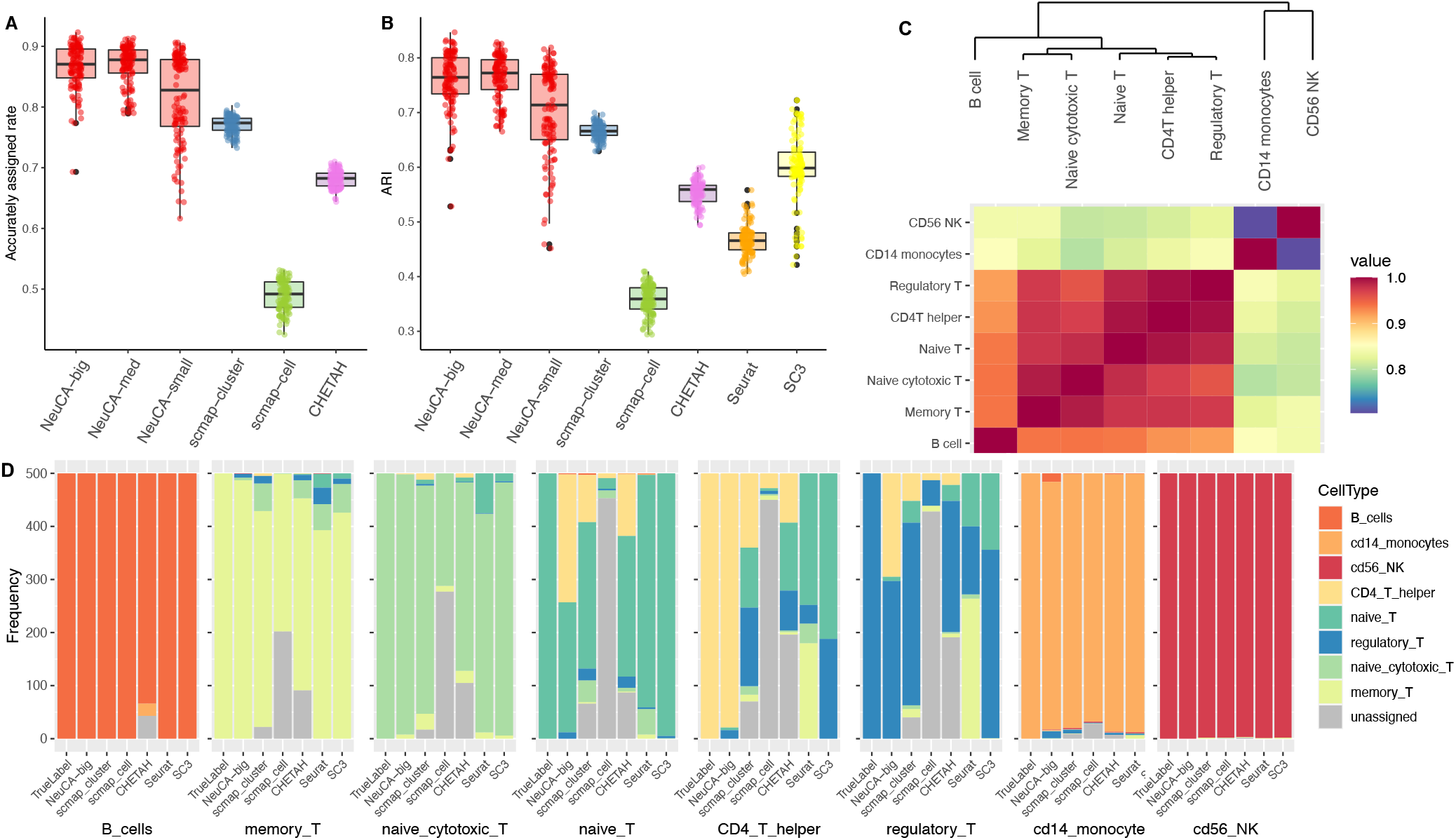
Cell classification accuracy results in the numerical study of the 10X PBMC dataset. The presented results were summarized over 160 Monte Carlo simulations. (A) and (B) show the accurately assigned rate and the adjusted Rand index (ARI) of the proposed and existing methods. (C) shows the hierarchical clustering results and correlations of the cell types. (D) is a detailed breakdown of the frequency counts of predicted labels using the proposed and existing methods, for each of the eight cell types, respectively.

We then explored in detail how NeuCA outperforms existing methods, and in what cell types the advantage is prominent. Figure 2C provides an illustration of the correlation and hierarchical structure of the cell types and Figure 2D presents a detailed breakdown of the frequency counts of the predicted labels from all of the methods. The true label is shown on the left-most bar in each sub-panel of Figure 2D. Combining Figure 2C and 2D, we found CD56 natural killer (NK) and CD14 monocytes are the two distinct cell types that are less correlated with other cell types. This makes them easy to be correctly predicted. As expected, all of the methods work well in annotating CD56 NK and CD14 monocytes (the last two sub-panels in Figure 2D). For closely correlated cell types, for example, naive cytotoxic T and memory T, existing methods annotate part of the cells as “unassigned” or incorrectly assign them cell labels. NeuCA demonstrates exceptional accuracy in annotating both naive cytotoxic T and memory T with few mistakes. Lastly, naive T, CD4 T helper, and regulatory T are the three most challenging cell types to predict, because the correlation between them is higher than 0.95. For these three cell types, existing methods produced more than 50% “unassigned” labels owing to uncertainty. In contrast, NeuCA can accurately annotate the majority of cells with a small portion of incorrect labels for these three cell types. These findings show that adopting hierarchical structure and step-specific feature selection improves the classification, especially in closely correlated cell types.

### Effect of training and testing sample sizes

One potential concern of adopting NeuCA is whether the model needs a large training dataset to achieve satisfactory performance. Therefore, we evaluated NeuCA and existing methods with different amounts of training data. Note that unsupervised methods Seurat and SC3 do not use training sets at all, and thus their performance stays similar over various settings. Supplementary Figure S1 shows the accuracy and ARI with training sizes ranging from 10%, 20%, 50%, and 80% of all cells. Training sets with 10% correspond to ∼6000 cells (∼750 cells per cell type), which is easy to achieve for real-world experiments. The testing dataset stays at 4000 randomly selected cells. These results show the high accuracy of NeuCA with only 10% cells used as the training data. NeuCA also consistently outperforms existing methods using 20% (and more) of cells for training. The accurately assigned rate increases from 0.86 to 0.90, which indicates the advantage of using a large training set of around 50,000 cells, although the improvement is marginal. From these experiments, we found that a training set with around 1000 cells per cell type is sufficient to provide a reasonably high annotation accuracy. In general, this is attainable in real-world single-cell RNA-seq experiments.

We also evaluated the impact of testing sample size on the performance of all methods. Supplementary Figure S2 reports the accurately assigned rate and ARI under different testing sample sizes (i.e., 800, 1600, and 4000 cells). As expected, changing the test sample size has little impact on the relative accuracy of supervised methods. The unsupervised methods have some performance improvements, with ARI increasing from 0.45 to 0.5 for Seurat and 0.60 to 0.65 for SC3. This also is within our expectation because increased sample size means more information for use. Nevertheless, NeuCA demonstrated a higher accuracy and ARI than the compared existing methods.

### Effect of highly and lowly correlated cell types

Last, we evaluated all of the methods under different prediction difficulty levels. As discussed in the Introduction section, the cell annotation becomes challenging when cell types are highly correlated. Using the PBMC data, we specifically designed two settings: one with highly correlated cell types only, and another one with lowly-correlated cell types only. These two scenarios correspond to difficult and easy prediction task, respectively.

For difficult prediction task, we only retained the five T-cell types from the PBMC data. We tested under different training and testing proportions as described in the section “Effect of training and testing sample sizes”. Supplementary Figure S3 shows an overall summary of the performance. All methods have decreased performance under this difficult setting. With the adoption of a hierarchical structure and step-specific feature selection, NeuCA still outperforms all existing methods in both accuracy and ARI. Interestingly, we find SC3 achieves the highest ARI over all existing methods. The high unassigned rate due to the high correlations of cell types lowers the accuracy and ARI of existing supervised methods. Supplementary Figures S4 and S5 demonstrate the desirable performance of NeuCA under different training and testing sizes. Similar conclusions can be drawn from the experiments in the easy prediction setting (Supplementary Figures S6 - S8). In this relatively easy cell labeling problem, all methods yield reasonable performance. Our method still leads the competing ones, with almost perfect accuracy on average even when using only 10% of the data to train the model.

### Applications on cell sorted human PBMC datasets

Next, we evaluated our proposed method, NeuCA, along with other existing methods, on three additional PBMC-related real datasets: PBMC_8ct_random500 is a subset of the aforementioned FACS-sorted PBMC data with 500 cells randomly drawn for each cell type for a total of 4000 cells^26^; Zhengmix_8ct_PBMC is also from the 10X platform^26^ and has been used in several benchmarking studies^27^; FACmix_NK_Mono is a two-component mixture of FACS-sorted NK cells^28^ and FACS-sorted monocytes^29^, with a total of 12,700 cells. Compared with the first two datasets that include eight distinct PBMC cell types, the third one only has two cell types. However, the third dataset is obtained from two independent studies, and thus valuable in evaluating the inter-study performance of all of the methods.

To evaluate the robustness of the proposed method, we trained NeuCA using one dataset and tested on another one, with various combinations of the datasets described above. The third dataset was not used as a training set due to smaller cell type number. The same training and testing strategy is adopted on other existing supervised methods (i.e., scmap-cluster, scmap-cell, and CHETAH) for benchmarking purposes. The results for all supervised methods are presented in Figure 3. Consistent with our findings in the previous numerical studies, NeuCA achieves more accurate label assignment than existing methods in intra-study predictions (the first and third panels in Figure 3A). NeuCA also has outstanding performance in correctly annotating the inter-study dataset and obtained more than 98% accuracy (second and fourth panels in Figure 3A). In comparison, the performance of existing supervised methods is less stable in the inter- and intra-study experiments. We found that scmap-cluster and CHETAH have good performance in the intra-study experiments, but they fare worse in the inter-study settings, while scmap-cell has a reverse pattern.

**Figure 3.**
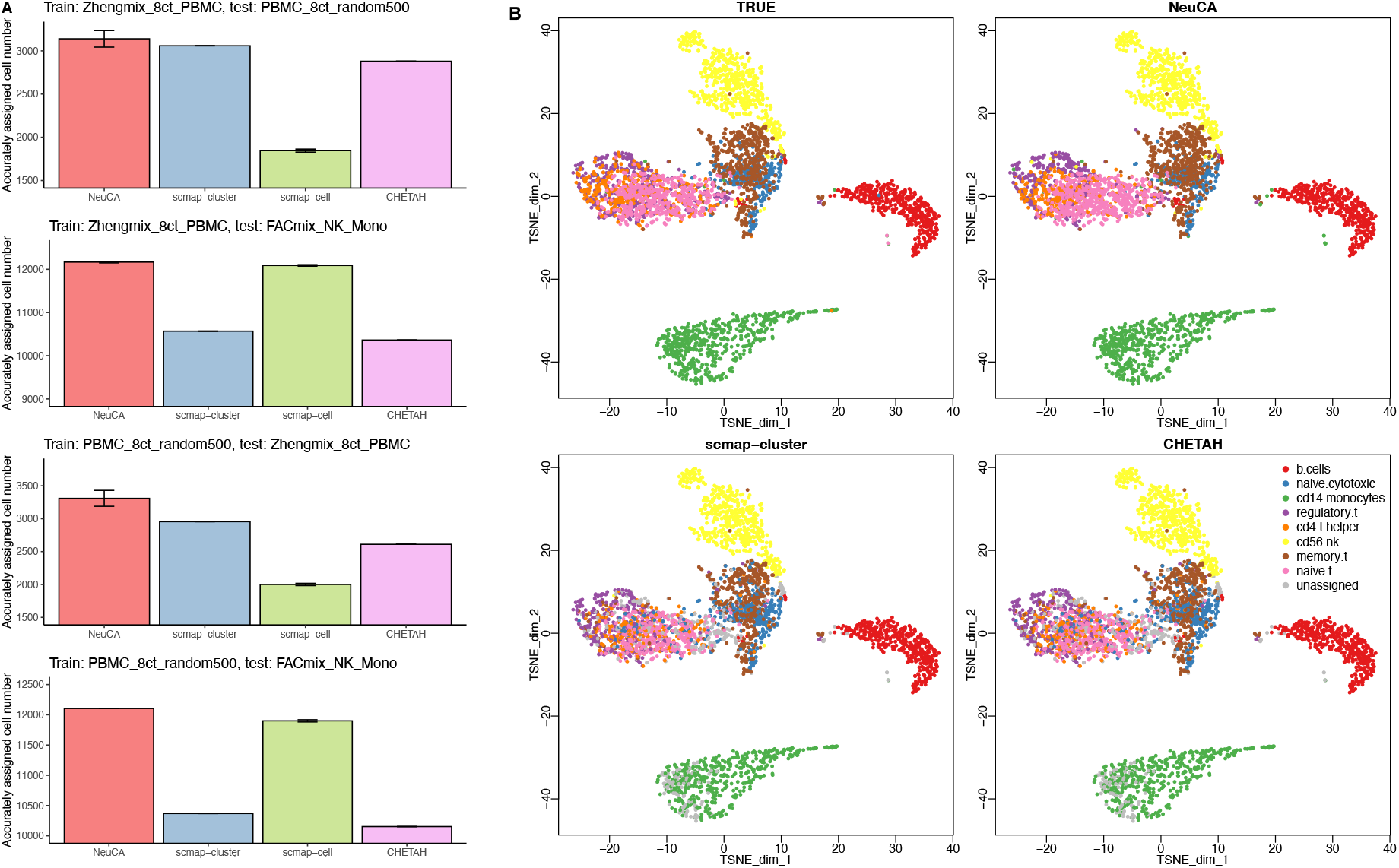
Results from applying NeuCA and existing methods on three PBMC datasets. (A) shows the accurately assigned cell number for different methods. (B) shows the *t*-SNE clustering plot of the cells, using *PBMC*_8*ct*_*random*500 as the training data and *Zhengmix*_8*ct*_*PBMC* as the testing data. Note NeuCA has high concordance with the ground truth, with very few unassigned cells (grey color cells).

We presented the low dimension t-distributed stochastic neighbor embedding (*t*-SNE) visualization of the true and predicted cell types for NeuCA, scmap-cluster, and CHETAH in Figure 3B to provide additional insights into the performance differences. The results are based on a specific scenario where *PBMC*_8*ct*_*random*500 was used as the training data and *Zhengmix*_8*ct*_*PBMC* as the testing data. Both scmap-cluster and CHETAH have a mixture of correct and incorrect predictions for the cloud of T cells with a considerable proportion of unassigned cells. In comparison, NeuCA has more similar patterns to the true labels in all major cell lineages. Additional visualizations for scmap-cell and the two unsupervised methods, Seurat and SC3, are presented in Supplementary Figure S9.

### Applications on human pancreas datasets

We applied NeuCA and other methods on four human pancreas datasets to specifically evaluate the inter-study classification performance. The four real datasets are: Baron^21^ data, Muraro^30^ data, Seg^31^ data, and Xin^32^ data. These four studies contained different numbers of cells, were obtained from different numbers of subjects, and were sequenced using different sequencing protocols. The number of subjects ranges from 4 to 18, and the number of cells ranges from 700 to >8,000. Baron, Muraro, Seg, and Xin used inDrop, CEL-Seq2, Smart-Seq2, and SMARTer, respectively. To comprehensively evaluate the inter-study annotation accuracy, we iteratively used each of the four as the training set and examined the performance in the remaining three datasets. We used the published cell labels as the gold standard annotations. To have true labels consistency in the four cell types, we processed the four datasets by first removing rare cell types. This allowed for inter-study comparisons. The four processed datasets had 1,600, 2,038, 8,569, and 2,126 cells in Xin, Seg, Baron, and Muraro datasets, respectively.

Figure 4A-D shows that NeuCA achieves the highest accuracy among 9 of the 12 scenarios, and comparable accuracies for the remainder. In several settings, the advantage of NeuCA is substantial compared with other methods. For example, when using Baron for training and Seg for testing, NeuCA accurately assigned labels for more than 90% of the cells, while the competing methods only annotated 80% or less cells correctly. The scmap-cell method was the second-best performer in general, which gave stable performance across the scenarios.

**Figure 4.**
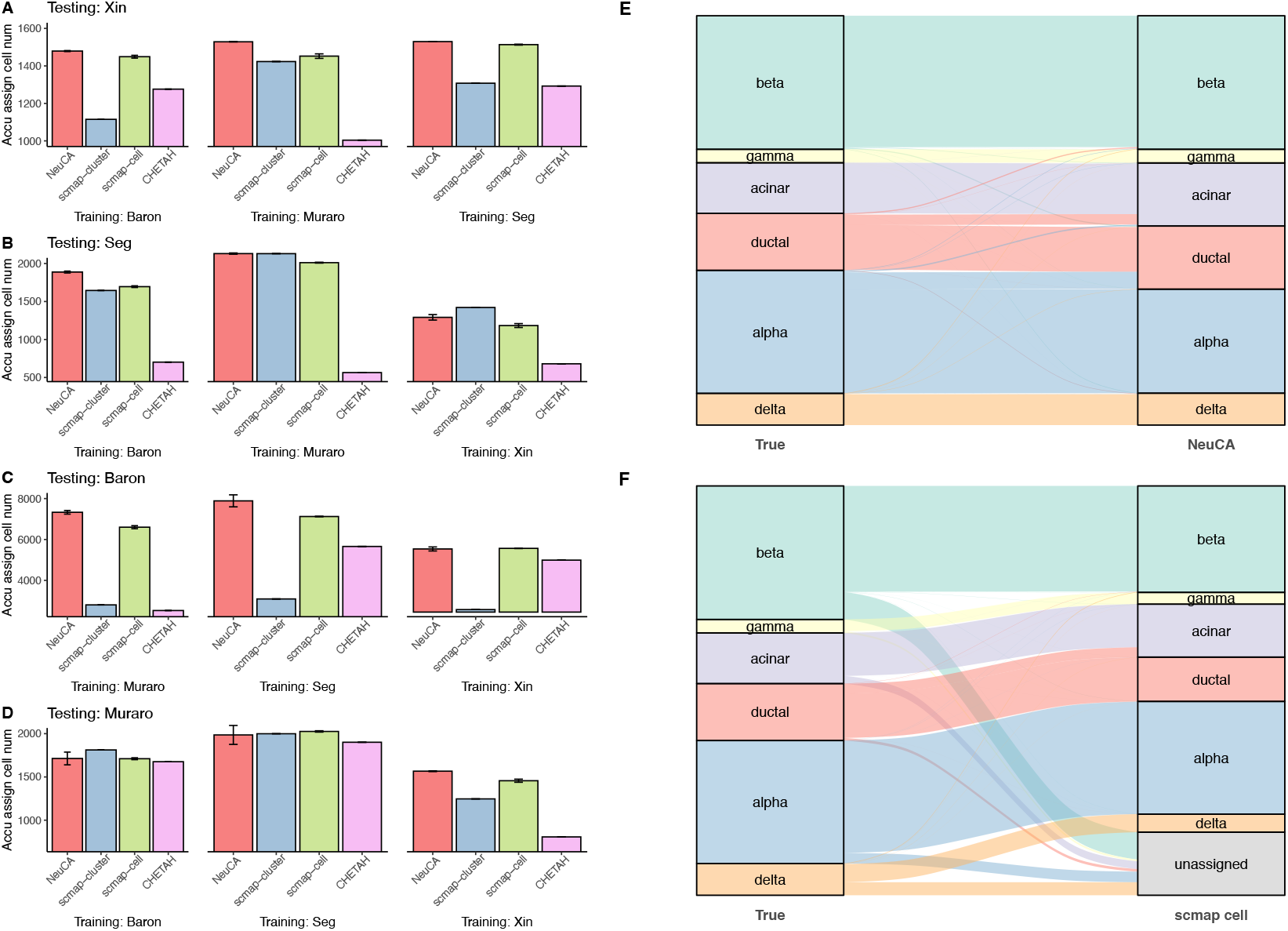
Cell label prediction results from applying the proposed method on four human pancreas datasets. The total cell numbers for the four datasets are 1,600, 2,038, 8,569, and 2,126. Panel (A)-(D) shows the accurately assigned cell numbers from proposed and existing methods, alternating training and testing dataset selections. Panel (E) and (F) show the comparison of true labels (left column) and the estimated labels (right column) from NeuCA and scmap-cell, using Sankey diagrams, with Seg data as the training and Baron data as the testing dataset.

Figure 4E and Figure 4F are detailed examinations of the classification results of NeuCA and scmap-cell using Sankey diagrams. In the diagram, the width of the flows reflects the frequency of cells in each cell type. For both diagrams, we trained the model using the Seg data and tested the model using Baron data. The box column on the left shows the true frequencies of cells, while the box column on the right shows the predicted frequencies of cells. A one-to-one mapping relationship is shown in the flow in-between. NeuCA has higher overall accuracy than scmap-cell. Although the relative proportions of NeuCA and scmap-cell are similar, scmap-cell suffers from a high proportion of unassigned cells. scmap-cluster and CHETAH also suffer from high proportions of unassigned cells, leading to lower accurately assigned cells numbers (Supplementary Figure S10). Similar conclusions can be drawn from Sankey diagrams of the unsupervised learning methods Seurat and SC3 (Supplementary Figure S11).

### Applications on ASD data

Lastly, we benchmarked all methods on a cross-condition annotation problem. We obtained a set of single nucleus RNA-seq data from a study for Autism spectrum disorder (ASD)^33^, a group of cognitive developmental disabilities that cause significant social, communication, and behavioral challenges^34^ for patients. This study contains snRNA-seq data from 15 ASD patients and 16 controls. Compared with the previous datasets, this is a much larger dataset with 52,003 cells in the ASD group and 52,556 cells in the control group. We were interested in evaluating the methods across the disease conditions, i.e., accurately predict cell types in the ASD group using the control data as the training set or vice versa. This evaluation is motivated by the pragmatic consideration that often the existing single cell data only contain normal samples, while researchers are interested in using the information in annotating the diseased subjects.

Figure 5 shows the accuracy and ARI for two scenarios: ASD samples as the training set and control samples as the testing set (Figure 5A); control samples as the training set and ASD samples as the testing set (Figure 5B). NeuCA achieves the highest accuracy and ARI among all methods. Interestingly, NeuCA has better performance in predicting control samples than predicting ASD samples. We suspect that ASD-related molecular changes might be associated with such performance change, e.g., the differentially expressed genes make it harder to accurately annotate cells in ASD samples. scmap-cell is the second-best method in the supervised category with an accurately assigned rate around 0.7. Unsupervised methods Seurat and SC3 also have good performance, due to the large number of cells. This experiment shows that NeuCA stably outperforms existing methods in cross-condition annotations.

**Figure 5.**
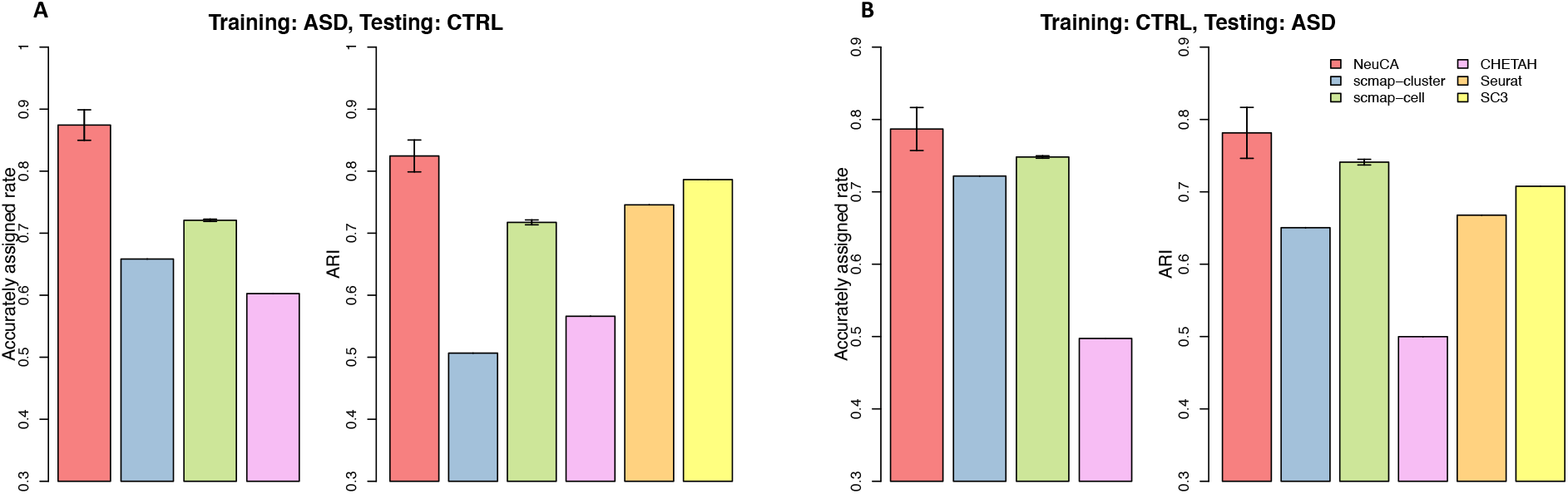
Accurately assigned rate and ARI of the proposed and existing methods on the ASD data, containing both ASD disease samples and control samples. Panel (A) shows the accuracies using ASD disease samples as the training set and control samples as the testing set. Panel (B) shows the results using control samples as the training set and ASD disease samples as the testing set. NeuCA stably outperforms existing methods in cross-condition annotation tasks.

## Discussion

We have developed a supervised learning method, NeuCA, for cell label assignment in single cell RNA-seq data analysis. Our method uses existing scRNA-seq data with known labels to train a neural network based classifier, adopting different strategies for highly correlated and average cell types. In situations where highly correlated cell types are absent, NeuCA adopts a feed-forward neural network for cell type classification. In situations where highly correlated cell types exist, NeuCA adjusts itself to a hierarchical neural network with an inferred cell lineage tree structure and stepwise feature selections. This flexible approach is accurate and efficient compared with existing methods, as shown in the series of numerical simulations and eight real data applications.

NeuCA aims to alleviate two problems in existing cell type annotation methods. First, data analysis using unsupervised clustering methods involves identifying cell type-specific markers and manually assigning labels. Such a step is usually laborious and time-consuming. Also, the lower dimensional projection of clusters does not necessarily represent the biologically-meaningful cell types. Moreover, manual labeling could be subjective and lack reproducibility. Second, many existing supervised approaches, such as scmap-cluster, scmap-cell, and CHETAH, fail to distinguish uncertain assignment from unknown cells, resulting in a high proportion of unassigned cells. When the cell types are well-studied, such as in PBMC and human pancreas tissues, these unassigned labels lead to an unwelcomed loss of cells in downstream analysis. Our investigation shows that existing methods tend to have difficulty in distinguishing highly correlated cell types, which leads to the unassignment problem. NeuCA’s flexible and efficient two-step approach offers solutions to both aspects.

It is worth noting that NeuCA is designed to fit the situations where cell types are well-studied, i.e., the situations where all the cell types in the new sample already exist in the reference/training data. This may appear to be very restrictive at first glance, but NeuCA is still widely applicable. First, as shown in our real data experiments, many tissue types have already been comprehensively sequenced and labeled in previous scRNA-seq studies. For example, the Human Cell Atlas (HCA) is on track to create reference maps of all human cells, providing a rich resource for obtaining comprehensive reference data. Currently HCA has curated more than 12 million cells from 55 human organs. Another example is the Mouse Cell Atlas (MCA), where over 1.2 million single cells from more than 28 mouse tissues are available. These collections provide rich resources for reference/training data for NeuCA. Second, when one expects the existence of unknown cell types in the collected samples, there are tools available for identifying those rare or unknown cells from the data^35,36^. This can be done as the first step prior to applying NeuCA. After unknown cells are identified, we can use NeuCA on the rest of the cells to assign labels. One example is with tumor samples, where alternative methods, such as inferCNV^4^ and copyKAT^37^, could be applied to identify malignant cells first and NeuCA at the second step. Third, it may not be easy to decide whether unknown or novel cell types exist in the data. Although our application to ASD data shows that NeuCA can be applied to cross-condition predictions, this may not hold in other scenarios. For example, previous studies have reported that certain types of hematopoietic stem and progenitor cells (HSPC) play an important role in myelodysplastic syndromes (MDS) and that the presence of HSPC subtypes varies considerably in MDS patients^38,39^. We recommend consulting biological/clinical experts to make related decisions when analyzing tissues from novel conditions.

We have implemented NeuCA in an R/Bioconductor package. NeuCA uses *keras* as the application programming interface (API) to enable fast high-level neural network implementation. The software builds and deploys TensorFlow within R programming, and it can run on either CPU or GPU devices. Additionally, we provide a trained classifier for human PBMC data that can be directly applied. Our proposed NeuCA method is available at https://bioconductor.org/packages/NeuCA/ with a detailed user manual document.

## Methods

Suppose the training data consist of scRNA-seq expression matrix **X**_0_ and the known cell type labels ***y***_0_. Each row of ***X*** _0_ is a gene and each column is a cell. ***y***_0_ is a vector with the same length as the total cell number (column number) in the training data. Suppose there are *K* distinct cell types. Given a testing scRNA-seq expression matrix **X**_1_, the goal of NeuCA is to predict the cell labels ***y***_1_ based on **X**_0_, **X**_1_, and ***y***_0_. After the preprocessing steps, such as normalizing the expression matrix and taking subsets of ***X***_0_ and ***X***_1_ by their overlapped genes, NeuCA will train a classifier using a feed-forward neural network or a hierarchical neural network, depending on the absence or presence of highly correlated cell types. The detailed steps are described next.

### Correlation and NeuCA strategy

NeuCA first computes the averaged expression profile matrix for all the cell types in the training data. This is a straightforward averaging step, using the mean expression values of the cells for each cell type. Next, with the cell type-specific averaged expression profiles available, we compute the correlation matrix, **P** = (*p*_*i, j*_), where *p*_*i, j*_ is the Pearson’s correlation coefficient of cell type *i* and *j*. In the cases where all the off-diagonal elements in ***P*** are smaller than a threshold of *τ*, NeuCA trains a feed-forward neural network using the training data and predicts the labels in the testing data. Otherwise, when highly correlated cell types exist, it uses a hierarchical neural network tree as the classifier. The threshold *τ* quantifies the cut-off level of correlations of highly correlated cell types. We used *τ* = 0.95 across all of the numerical simulations and real data experiments.

### Feed-forward neural network

Given the training scRNA-seq matrix ***X*** _0_ and the cell label vector ***Y***_0_, the feed-forward neural network with *L* hidden layers has a standard architecture

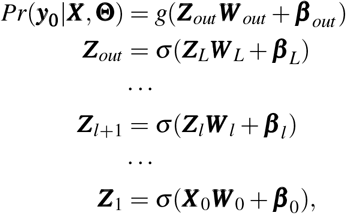

where **Θ** ={***W*** _0_, … ***W*** _*l*_, … ***W*** _*L*_, ***β*** _0_, … ***β*** _*l*_, … ***β*** _*L*_, ***W*** _*out*_, ***β*** _*out*_}, ***Z***_*l*_, for *l* = 1, … *L* − 1, are the hidden neurons with corresponding weight matrices *W*_*l*_ and bias vectors ***β*** _*l*_. The dimensions of ***Z*** and ***W*** depend on the number of hidden neurons, the input dimension, and the number of cell types *K. σ* (·) is the activation function such as sigmoid, hyperbolic tangent, or rectifiers. *g*(·) is the softmax function that converts the values of the output layer into predicted probabilities. For example, for the *i*-th cell in the training set,

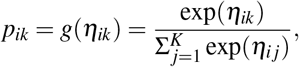

where *p*_*ik*_ = Pr(*y*_*i*_ = *k*|***x***_*i*_), 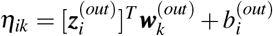 and *i* = 1, …, *n*. The parameters to be estimated in the feed-forward neural network are all the weights and biases, i.e., **Θ**. We train the model using a stochastic gradient descent (SGD) based algorithm with the following categorical cross-entropy loss as the objective function:

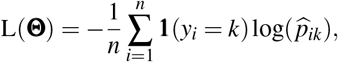

**1**(·) is an indicator function, and equals 1 only when the expression inside the bracket holds.

The activation functions used here were the rectified linear unit (ReLU), *σ*_*ReLU*_ (*z*) = *max*(*z*, 0), for a scaler variable *z*. The ReLU activation function has become the default activation for many types of neural networks because the model using ReLU is generally easier to train and often achieves better performance than alternative functions^40^. In addition, ReLU can avoid the vanishing gradient problem during optimization, which is an advantage over sigmoid and hyperbolic tangent functions^41^. For the optimization algorithm, we chose to use the RMSprop optimizer, as a recent comparison study shows the top accuracy using RMSprop over Momentem or AdaGrad, among others^42^. We also used the mini-batch training strategy that randomly trains a small proportion of the samples in each iteration^42^. We additionally have tried applying recent deep learning techniques on the training feed-forward neural network, such as batch normalization and dropout, and found the performance stayed similar with and without these techniques (data not shown).

In neural networks, the numbers of layers and nodes are also tunable parameters. Instead of changing these parameters in each and every application, we stratified them as three sets of choices in our experiments. We used 2, 3, and 4 hidden layers, respectively, for small, medium, and big model sizes. The number of nodes in the hidden layers are provided in Table 1. Although these choices may not be optimal over the full sets of parameter combinations, we found the current model setup can generate good performance in practice.

**Table 1.** Three different model sizes that can be selected in the feed-forward neural network classifier training.

| Neural network model size | No. of layers | No. of nodes in the hidden layers |
| --- | --- | --- |
| Small | 3 | 64 |
| Medium | 4 | 64, 128 |
| Big | 5 | 64, 128, 256 |

### Cell lineage-based hierarchical neural network

Our cell lineage-based hierarchical neural network is specially designed to handle the situation where highly correlated cell types exist. The training stage consists of four major steps. First, we construct a cell lineage hierarchical tree based on the average cell type profiles. This hierarchical tree will be used in the following step as a road map. In the second step, we label the regular correlated cell types using highly sensitive markers. We compute the highly sensitive marker genes for cell type *k* by

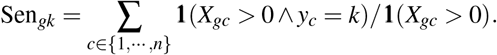

The genes with Sen_*gk*_ *>* 0.95 for each cell type were deemed as the cell type specific genes. For cell types with more than *v*_1_ highly sensitive markers, we identify the cells expressing more than *v*_2_ cell type-specific genes and assigned the corresponding cell labels. We use *v*_1_ = 10 and *v*_2_ = 3 for all of our experiments. We find that highly sensitive markers alone are already accurate in labeling regular cell types, but they cannot address highly correlated cell types. Third, we iterate the rest of the cells through the hierarchical tree and performed additional feature selection. At each bifurcation, we perform the differential analysis provided in limma^43^ to compare the cells from the left and right branches and select the top *M* genes as the bifurcation-specific features. We use *M* = 1000 in all of our experiments. Finally, at each bifurcation along the hierarchical tree, we train a feed-forward neural network to classify the cells from the upstream to binary outcomes (“left” or “right”). In each bifurcation node, we use the same feed-forward neural network setting and training as described in the “Feed-forward neural network” section above. The only difference is that we replace the categorical cross-entropy by a binary cross-entropy, defined as

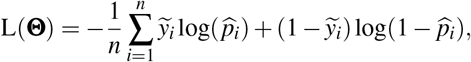

where 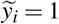 if *y* belongs to the right branch and 0 otherwise. When the classification reaches the output layer, the corresponding cell labels are assigned.

### Numerical simulations using real PBMC data

Real PBMC data from the 10X Genomics website (https://support.10xgenomics.com/)^26^ were downloaded and processed to generate our simulation datasets. The cells in the real PBMC data were FACS-sorted and consisted of 9,139 CD4 T helper cells, 7,959 B cells, 718 CD14 monocytes, 7,601 natural killer cells, 9,338 memory T cells, 10,993 naive cytotoxic T cells, 7,700 naive T cells, and 7,861 regulatory T cells. We designed three simulation settings to evaluate the methods under different scenarios. In the first setting, we randomly drew *p* proportion of cells as the training set. Among the rest of 1 − *p* proportion of cells, we drew *n* cells as the testing set. We consider *p* = 10%, 20%, 50%, 80% and *n* = 800, 1600, 4000. In the second setting, we focused the experiment on the T cells only. This aims to evaluate the performance of methods when all cell types are moderately or highly correlated. Thus, the five T cell types (i.e., CD4 T helper, memory T, naive cytotoxic T, naive T, and regulatory T) were considered here. In the third setting, we considered the major cell types and merged all of the five T cell types into one cell type: “T cells”. We used the same set of *p* and *n* in the second and third settings to evaluate the impacts of varying training and testing sample sizes.

### Real data analysis

In addition to the PBMC data used in the numerical simulation, we obtained another eight real datasets to validate the methods. The Zheng_8ct_PBMC data were downloaded from the R/Bioconductor package *DuoClustering2018*. The FACS sorted natural killer dataset was originally sequenced by^28^ and downloaded from the Gene Expression Omnibus (GEO, accession number: GSE130430). The FACS-sorted monocytes dataset was sequenced by^29^ and downloaded from GEO (GSE103544). The Xin^32^, Baron^21^, and Muraro^30^ datasets were also downloaded from GEO with accession GSE81608, GSE84133, and GSE85241, respectively. The Seg dataset^31^ was downloaded from the European Molecular Biology Laboratory (EMBL-EBI, https://www.ebi.ac.uk/, accession number E-MTAB-5061). Published cell labels were obtained for each study and used as true cell type labels.

## Acknowledgements

This study was supported by intramural funding from The University of Texas MD Anderson Cancer Center and Case Western Reserve University.

## Author contributions statement

Z.L. and H.F. conceived the models, conducted simulation and real data experiments, and wrote and reviewed the manuscript. H.F. developed the software package.

## Additional information

The authors declare no competing interests.

