## Supplementary Figure for "A neural network-based method for exhaustive cell label assignment using single cell RNA-seq data"

**Numerical experiments with cell-sorted PBMC data**
Supplementary Figure 1 and 2 show NeuCA's performance under various proportions of training and testing scenarios.

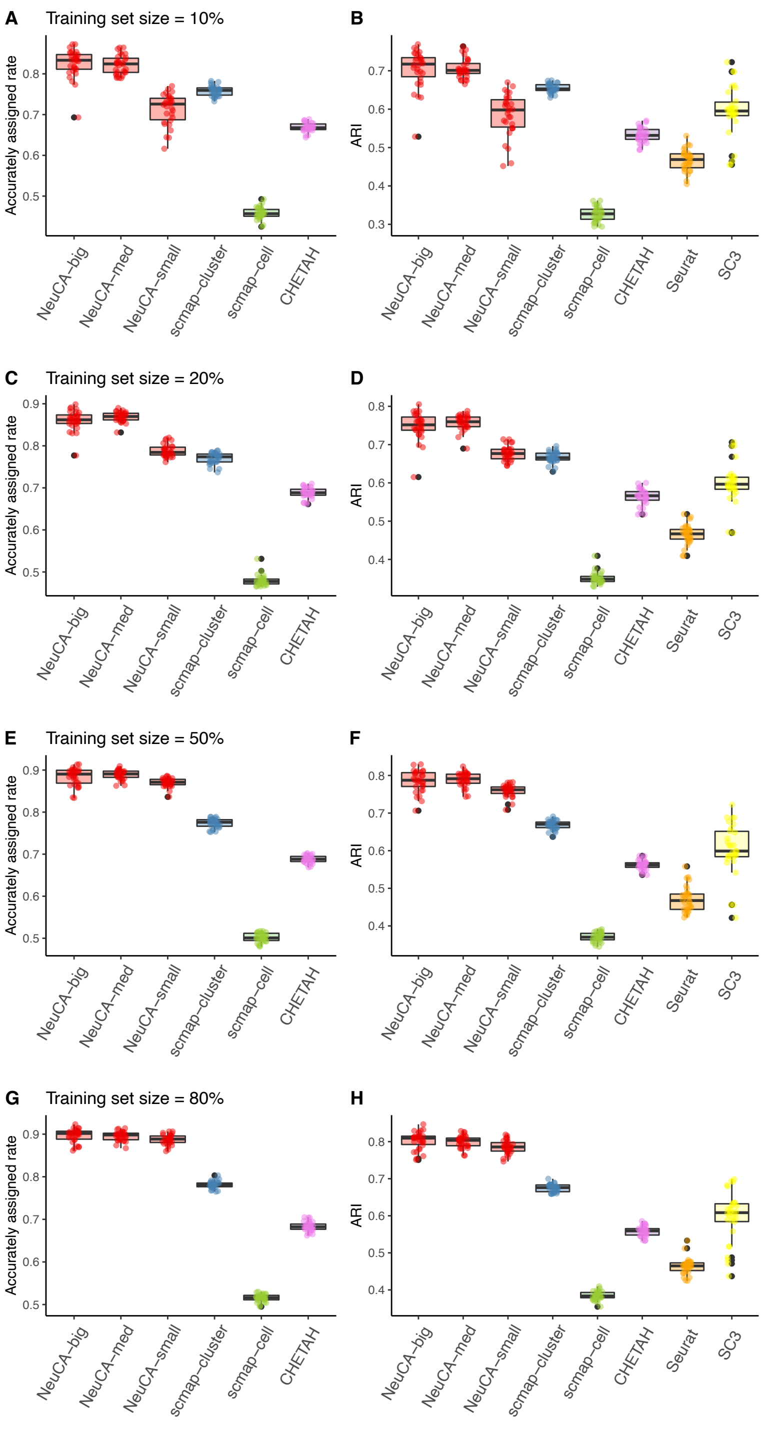


Supplementary Figure 1: Accurately assigned rate and ARI value using PBMC data with 8 cell types and different training set sizes. From top to bottom, the training set size increases from 10%, 20%, 50%, to 80%. Results are summarized over different testing size with each setting consisting of 20 Monte Carlo simulations.

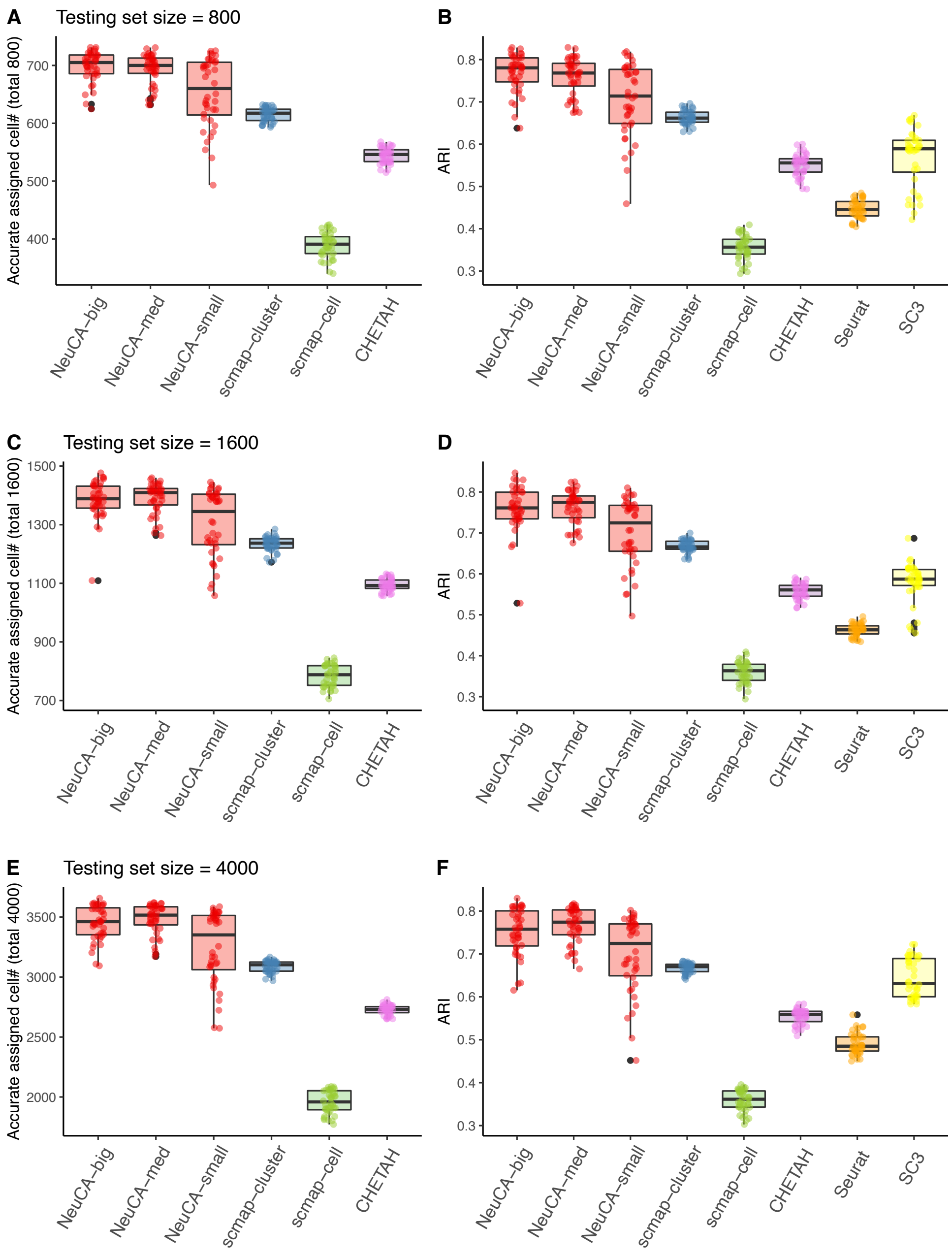


Supplementary Figure 2: Accurately assigned rate and ARI value using PBMC data with 8 cell types and different testing set size. From top to bottom, the testing set size increases from 800, 1600, to 4000. Results are summarized over different training size with each setting consisting of 20 Monte Carlo simulations.

**Numerical experiments exclusively for T cells**

Supplementary Figure 3 to 5 show NeuCA's performance at various split sizes of training and testing datasets, within hard-to-distinguish T cell subtypes.

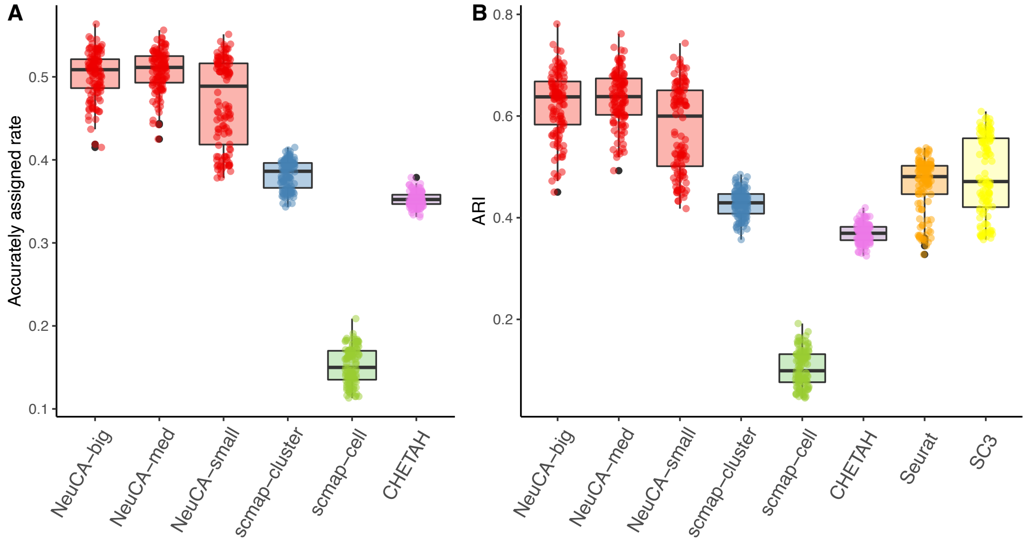


Supplementary Figure 3: Accurately assigned rate and ARI for applying the proposed method and existing methods on T-cell only dataset. Different training set sizes and testing set sizes are considered and summarized into one box for each method. 20 Monte Carlo simulations are used for each training/testing setting.

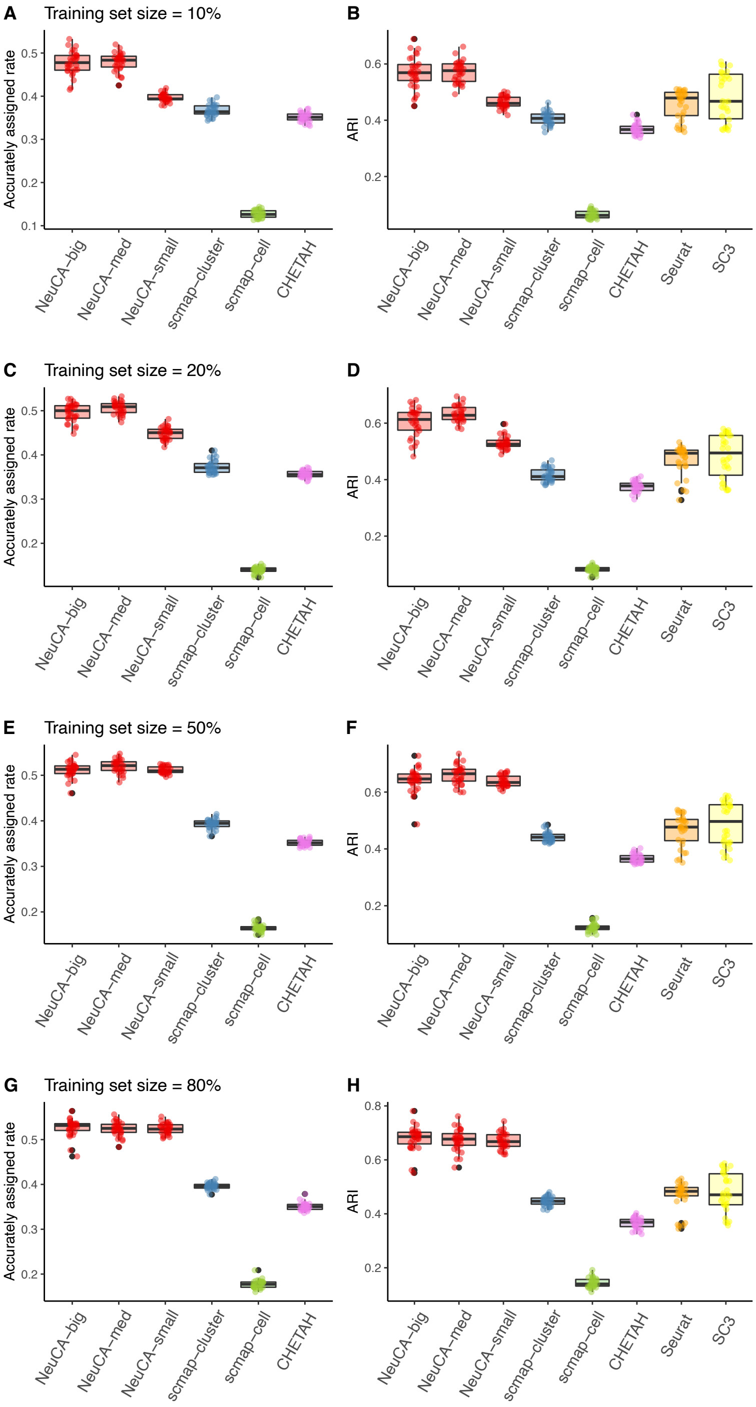


Supplementary Figure 4: Accurately assigned rate and ARI value using T-cell only data and different training set sizes. From top to bottom, the training set size increases from 10%, 20%, 50%, to 80%. Results are summarized over different testing size with each setting consisting of 20 Monte Carlo simulations.

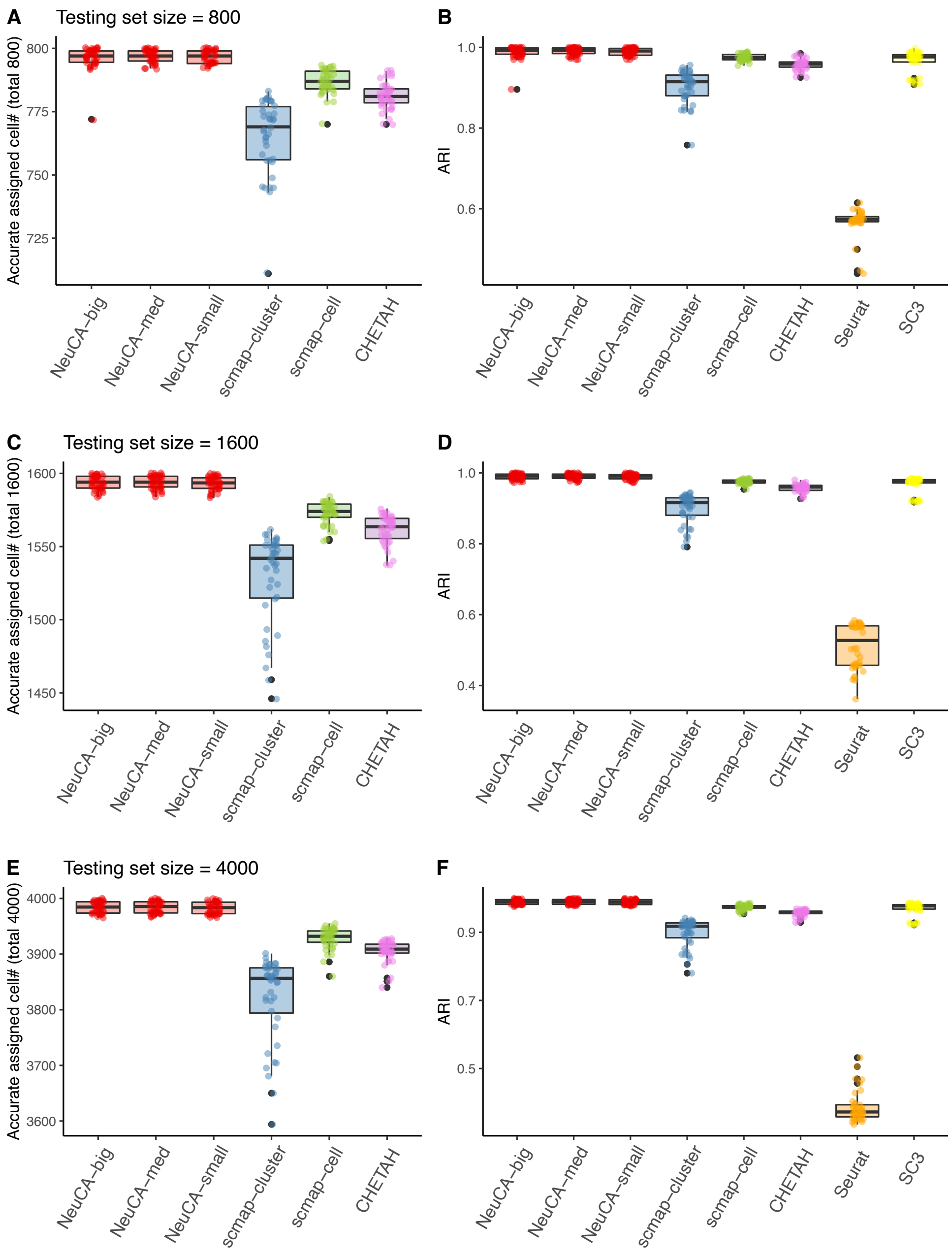


Supplementary Figure 5: Accurately assigned rate and ARI value using T-cell only data and different testing set size. From top to bottom, the testing set size increases from 800, 1600, to 4000. Results are summarized over different training size with each setting consisting of 20 Monte Carlo simulations.

**Numerical experiments exclusively for "easy" PBMC dataset**

Supplementary Figure 6 to 8 show NeuCA's performance for "easy" PBMC dataset. Here, all closely-correlated T cells were excluded from PBMC dataset, leading to very distinct cell types in the dataset that are easy to classify.

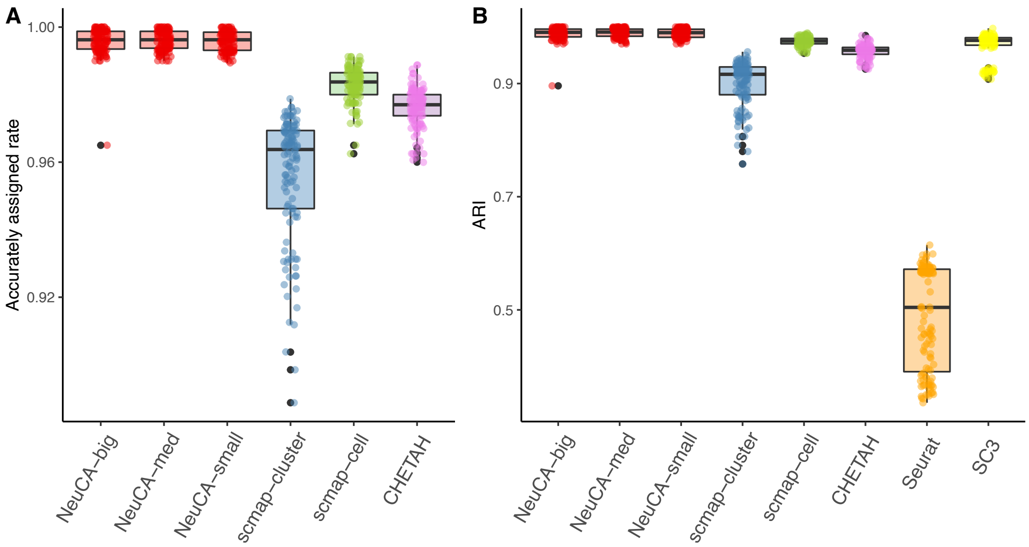


Supplementary Figure 6: Accurately assigned rate and ARI for applying the proposed method and existing methods on easy-PBMC dataset. Different training set sizes and testing set sizes are considered and summarized into one box for each method. 20 Monte Carlo simulations are used for each training/testing setting.

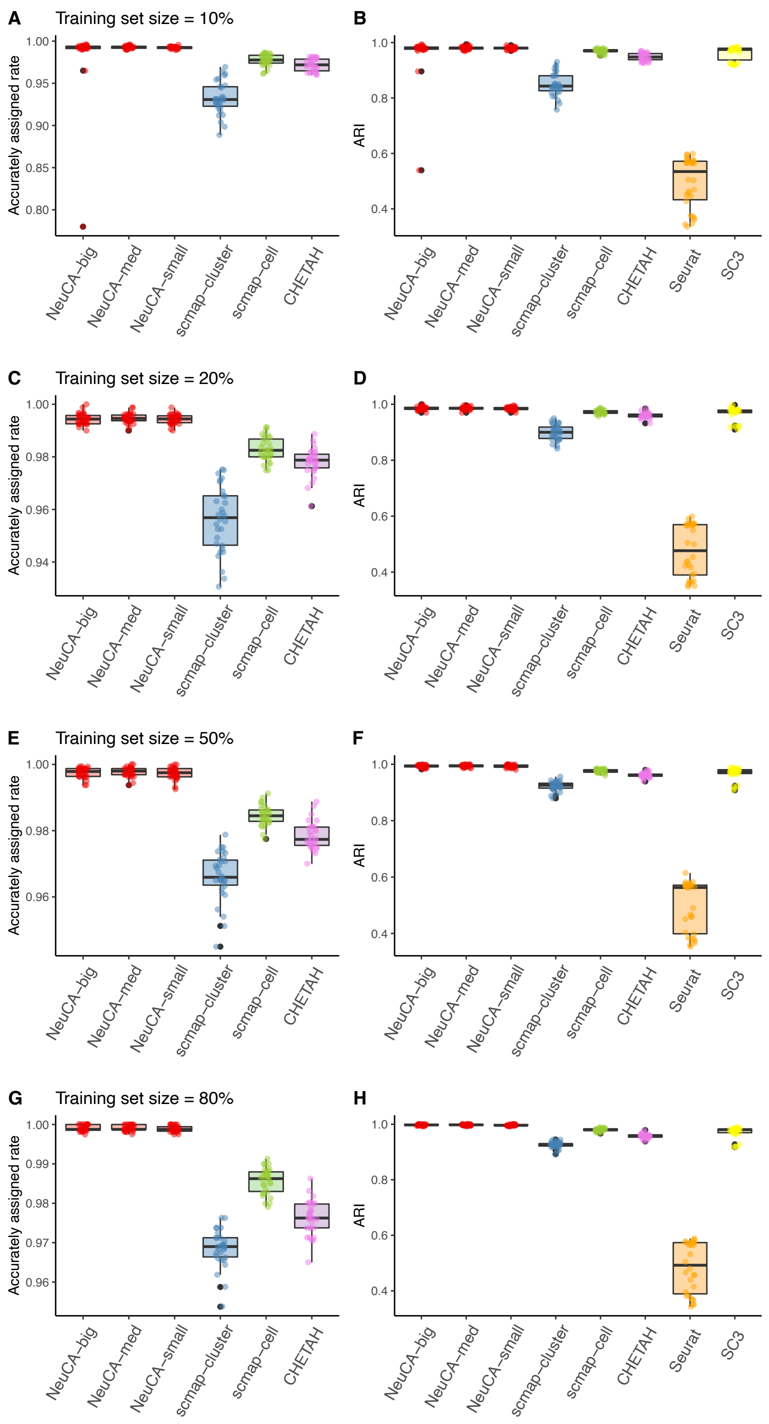


Supplementary Figure 7: Accurately assigned rate and ARI value using easy-PBMC data and different training set sizes. From top to bottom, the training set size increases from 10%, 20%, 50%, to 80%. Results are summarized over different testing size with each setting consisting of 20 Monte Carlo simulations.

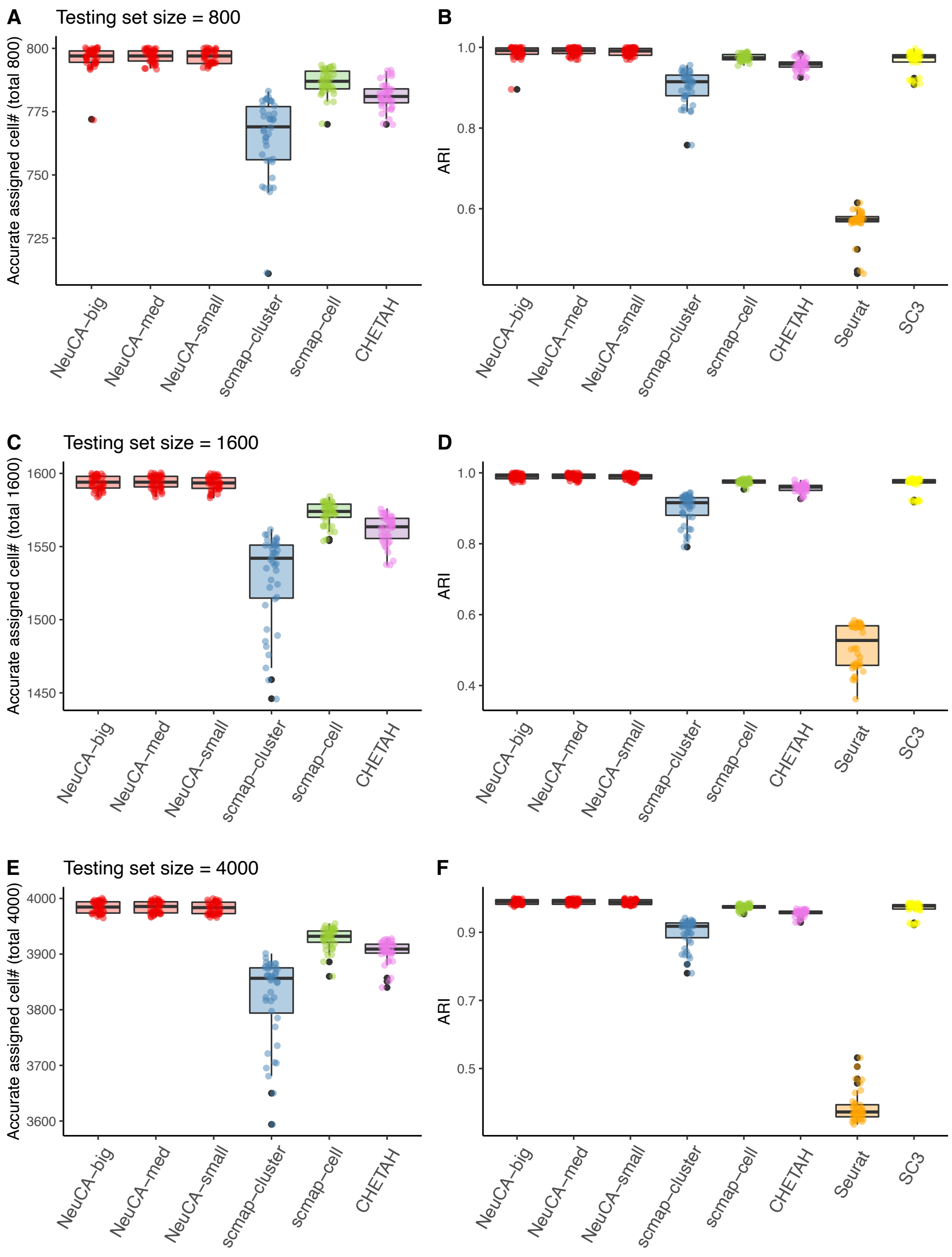


Supplementary Figure 8: Accurately assigned rate and ARI value using easy-PBMC data and different testing set size. From top to bottom, the testing set size increases from 800, 1600, to 4000. Results are summarized over different training size with each setting consisting of 20 Monte Carlo simulations.

**Real data analysis with cell-sorted PBMC data**

Supplementary Figure 9 is complementary to main Figure 3. It has additional visualizations of cell type annotating and clustering results using scmap-cell, Seurat and SC3. It is the result on real cell-sorted PBMC data.

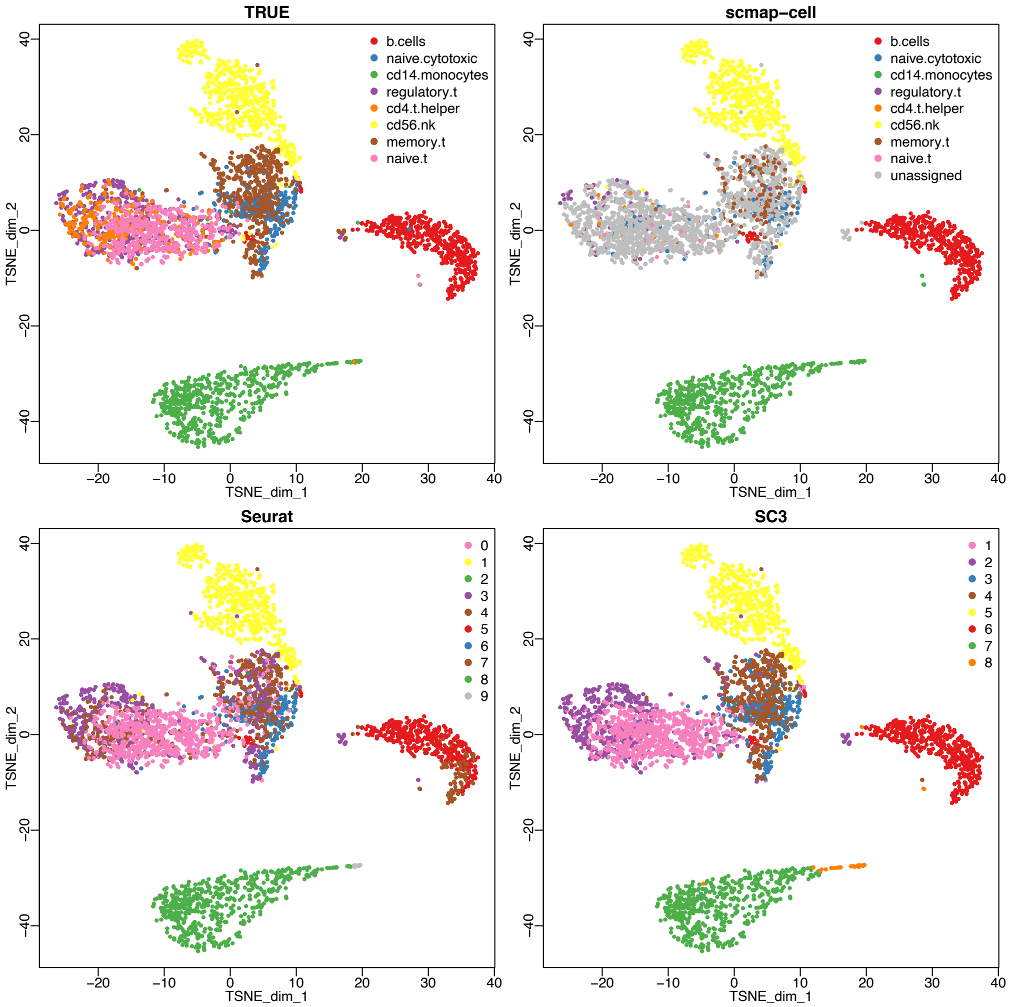


Supplementary Figure 9: Cell type annotating and clustering results from the PBMC real data experiment using scmap-cell, Seurat and SC3.

**Real pancreas data analysis**
Supplementary Figure 10 and 11 show Sankey diagram of predicted cell labels of 4 different methods.

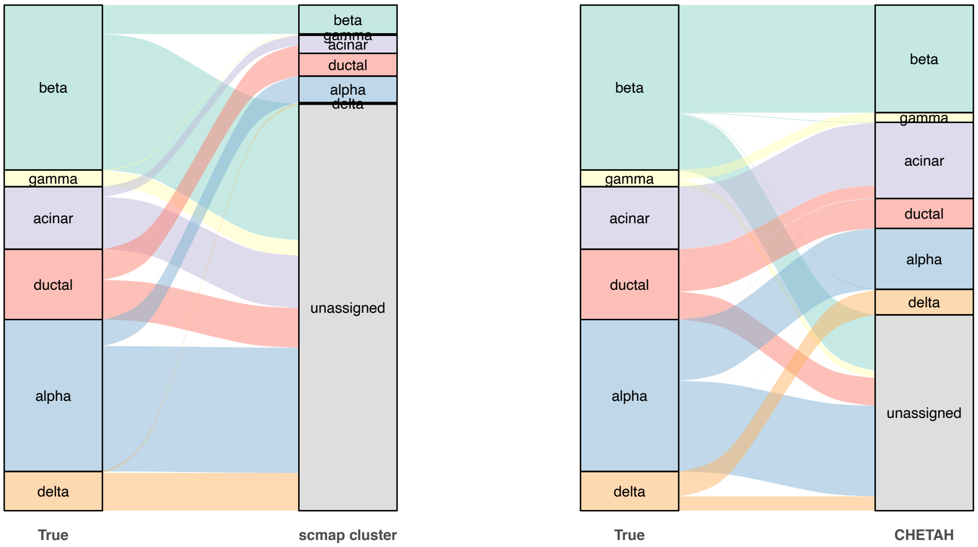


Supplementary Figure 10: Sankey plot of the true label and the estimated labels from pancreas data using scmap cluster and CHETAH. Seg data was used as training set and Baron data was used as testing set.

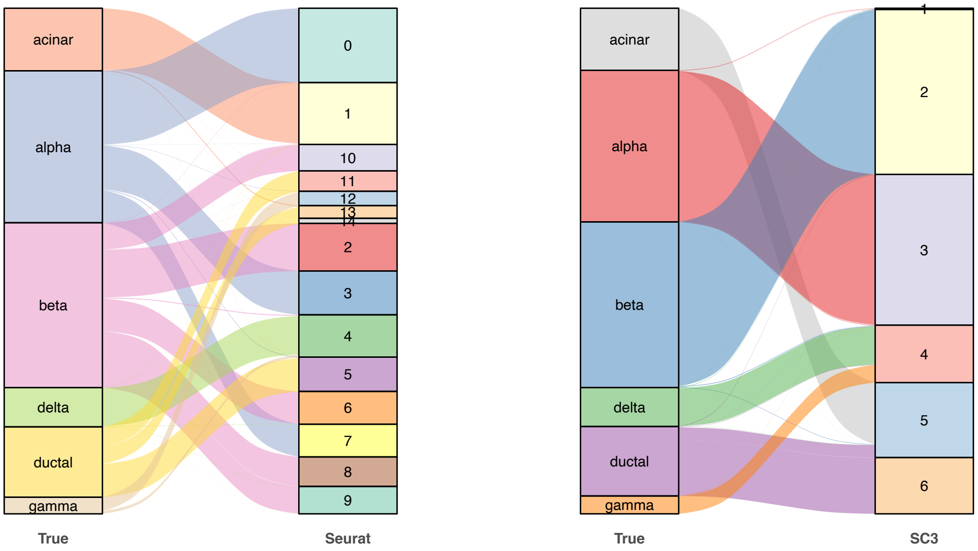


Supplementary Figure 11: Sankey plot of the true label and the estimated labels from pancreas data using Seurat and SC3. Seg data was used as training set and Baron data was used as testing set.
